# Metastasis is altered through multiple processes regulated by the E2F1 transcription factor

**DOI:** 10.1101/858068

**Authors:** Matthew Swiatnicki, Eran R. Andrechek

## Abstract

The E2F family of transcription factors is important for many cellular processes, from their canonical role in cell cycle regulation to other roles in angiogenesis and metastasis. Alteration of the Rb/E2F pathway occurs in various forms of cancer, including breast cancer. E2F1 ablation has been shown to decrease metastasis in MMTV-Neu and MMTV-PyMT transgenic mouse models of breast cancer. Here we take a bioinformatic approach to determine the E2F1 regulated genomic alterations involved in the metastatic cascade, in both Neu and PyMT models. Through gene expression analysis, we reveal few transcriptome changes in non-metastatic E2F1^−/−^ tumors relative to transgenic tumor controls. However investigation of these models through whole genome sequencing found numerous differences between the models, including differences in the proposed tumor etiology between E2F1^−/−^ and E2F1^+/+^ tumors induced by Neu or PyMT. Investigating mutated genes through gene set analysis also found a significant number of genes mutated in the cell adhesion pathway in E2F1^−/−^ tumors, indicating this may be a route for disruption of metastasis in E2F1^−/−^ tumors. Overall, these findings illustrate the complicated nature of uncovering drivers of the metastatic process.

## Introduction

Breast cancer is the most diagnosed cancer, and second leading cause of cancer death in women. To study underlying genomic events contributing to breast cancer, numerous genetically engineered mouse models have been generated, including the MMTV-Neu model [1] which recapitulates HER2+ve breast cancer, and the MMTV-Polyoma virus Middle T antigen (PyMT) [2] model. The PyMT model is highly aggressive, with tumors appearing as early as 45 days of age, and metastasis to the lung occurring in over 90% of tumor bearing mice. These phenotypes make PyMT a widely used model for the study of metastatic breast cancer. Similar to human breast cancers, both the Neu and PyMT models have striking heterogeneity at a histological, and gene expression level [3–6]. This heterogeneity reinforces the importance of these models as tools for the study of breast cancer.

Previous bioinformatic predictions using the Neu and PyMT models suggested a key role for the E2F1 transcription factor in Neu and PyMT tumors, suggesting that mechanisms outside the overexpression of the Neu or PyMT oncogene were contributing to tumor biology [7, 8]. The E2F family of transcription factors is involved in many important cellular processes, including apoptosis and cell cycle control. Usually sequestered by the retinoblastoma (Rb) protein, E2F1 is released to act on downstream targets once Rb becomes phosphorylated by cyclin dependent kinases (CDK) [9]. While mutations in E2F1 do not occur frequently in human breast cancer, mutations within the E2F pathway (Cyclin dependent Kinases, Retinoblastoma, etc.) occur in over 25% of breast cancer patients, illustrating the importance of the pathway [10–14].

To test the hypothesis that E2F1 regulated key events in Neu and PyMT tumors, E2F1 knockout (KO) mice [13] were interbred with Neu and PyMT models [7, 8]. This resulted in mammary tumors with altered phenotypic characteristics, including changes in latency, growth rate, and a significant decrease in metastasis to the lung. Metastasis is the ultimate cause of mortality in cancer, with an estimated 90% of cancer deaths resulting from the spread of cancer cells to distal sites within the body [15]. Typically, cancer cells undergo numerous important steps for completion of the metastatic cascade. These include escape from the primary tumor, intravasation, extravasation, and seeding the distal site [16]. The complicated nature of metastasis is illustrated in a recent review by Welch *et al* [17].

An important component contributing to the metastatic capability of a tumor is its microenvironment. Various collagens and proteins integral to cellular and tissue structure are capable of impacting metastatic potential. In fact, proteins within the extracellular matrix, including collagen IV, have been found to regulate metastasis within the liver [18]. Collagen IV is a major component of the basement membrane, which serves as an important barrier to tumor invasion. Various other proteins including lamanins, integrins, and fibronectin proteins are essential components of the extracellular matrix. The integrity of extracellular matrix, including the basement membrane, have been shown as being important for the early steps of tumor invasion and metastasis [19, 20]. Interestingly, a previous report demonstrated a decrease in the number of circulating tumor cells within PyMT E2F1^−/−^ mice, suggesting a disruption to the early steps in the metastatic cascade. Other data shows remodeling of the extracellular matrix at pre-metastatic lesion sites to be important for eventual seeding of distant metastasis [21].

Recent advances in bioinformatics methods have facilitated the investigation of cancer biology. Microarray and RNA sequencing studies have led to an abundance of publically available datasets, which allow for transcriptomic comparisons between primary tumor and distant metastatic lesions [22, 23]. Next generation sequencing has become widely available, and the sequencing of primary tumors as well as metastatic lesions has greatly increased our understanding of cancer genomics. Studies involving the sequencing of human tumors have described the mutation rate of solid tumors [24], as well as shown numerous genomic events are required for metastatic capability [25–27]. To study the underlying genomic events behind altered phenotypic characteristics in E2F1 KO tumors, whole genome sequencing was completed on E2F wild type and E2F1 knockout mammary tumors from the Neu and PyMT models. Here, we characterize the genome landscape of E2F WT and E2F1 KO tumors from both the Neu and PyMT models, and uncover new targets that may be critical to tumor development and progression.

## Results

We have previously demonstrated altered phenotypic characteristics upon ablation of E2F1 within the Neu and PyMT models. These included changes in growth rate and tumor latency upon interbreeding Neu and PyMT mice with E2F1^−/−^ mice (Figure 1A). Surprisingly, given the already shortened latency of PyMT mice, tumor latency in PyMT mice was significantly decreased upon E2F1 loss while growth rate remained unaffected. Interestingly, the opposite effect was seen within Neu E2F1^−/−^ mice, where latency was significantly increased, and growth rate was significantly decreased. Perhaps the most striking phenotype was a significant reduction of metastasis to the lung with loss of E2F1 in both mouse model strains (Figure 1B and C).

**Figure 1:**
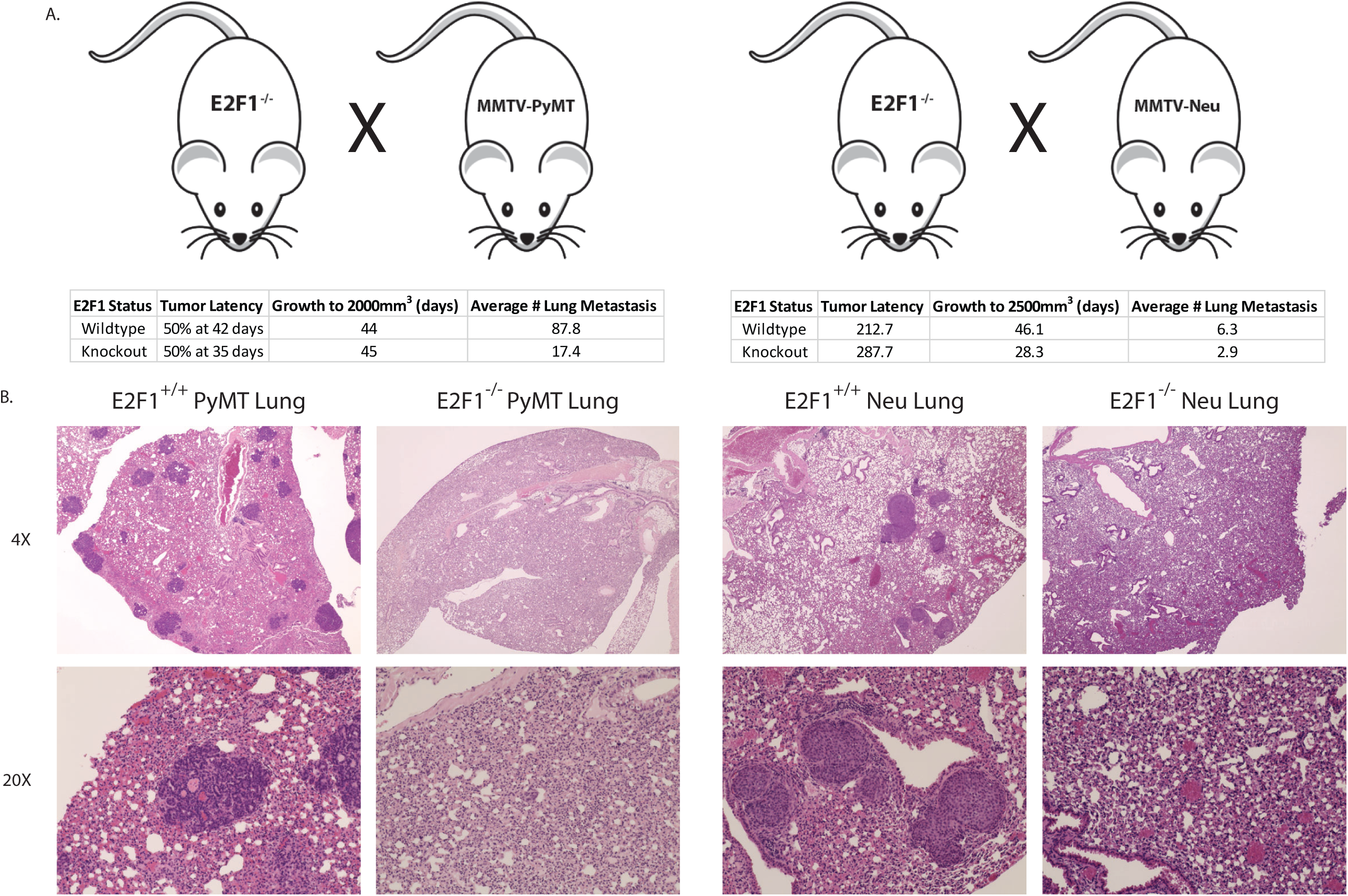
Altered phenotypic characteristics in E2F1^−/−^ tumors. **A)** E2F1^−/−^ mice were crossed with MMTV-Neu and MMTV-PyMT mice on the FVB background to create E2F1 knockouts in both models. **B)** Phenotypic changes seen in PyMT E2F1^−/−^ mice and (**C**) Neu E2F1^−/−^ mice, summarizing changes in latency, growth rate, and number of metastasis. H&E staining of E2F1^+/+^ mouse lung shows a large number of metastasis, while E2F1^−/−^ mice have little to no metastasis. Histology of the lungs was obtained at primary tumor endpoint.

To determine whether gene expression differences were responsible for phenotypic changes in E2F1 knockout tumors, microarray data were analyzed for fold change differences. Volcano plots revealed relatively few genes with major changes in gene expression data when analyzing equal to or greater than log2 fold changes between E2F1 WT and E2F1 KO tumors (Figure 2A). To elucidate whether this is recapitulated in human breast cancer, data from The Cancer Genome Atlas (TCGA) was analyzed. Microarray data from HER2+ve samples were isolated and E2F1 activity for these tumors was determined using pathway signature analysis. These samples were then stratified into quartiles for E2F1 activity and a log 2-fold change between genes was determined by subtracting samples in the lower quartile, from samples in the upper quartile. As shown by the volcano plot in figure 2B, human breast tumors resemble mouse mammary tumors in that low E2F1 activity does not lead to vast gene expression changes. To test for genetic pathways affected by loss of E2F1, Gene Set Enrichment Analysis (GSEA) was applied to expression data from Neu and PyMT E2F1 WT tumors in comparison to Neu and PyMT E2F1 KO tumors. GSEA analysis resulted in several differentially regulated pathways, including those involved in WNT signaling, and nucleotide excision repair (Figure 2C). This analysis highlighted the difficulties associated with the identification of pathways that were directly involved in breast cancer metastasis in bulk tumor samples from transcriptomic data.

**Figure 2:**
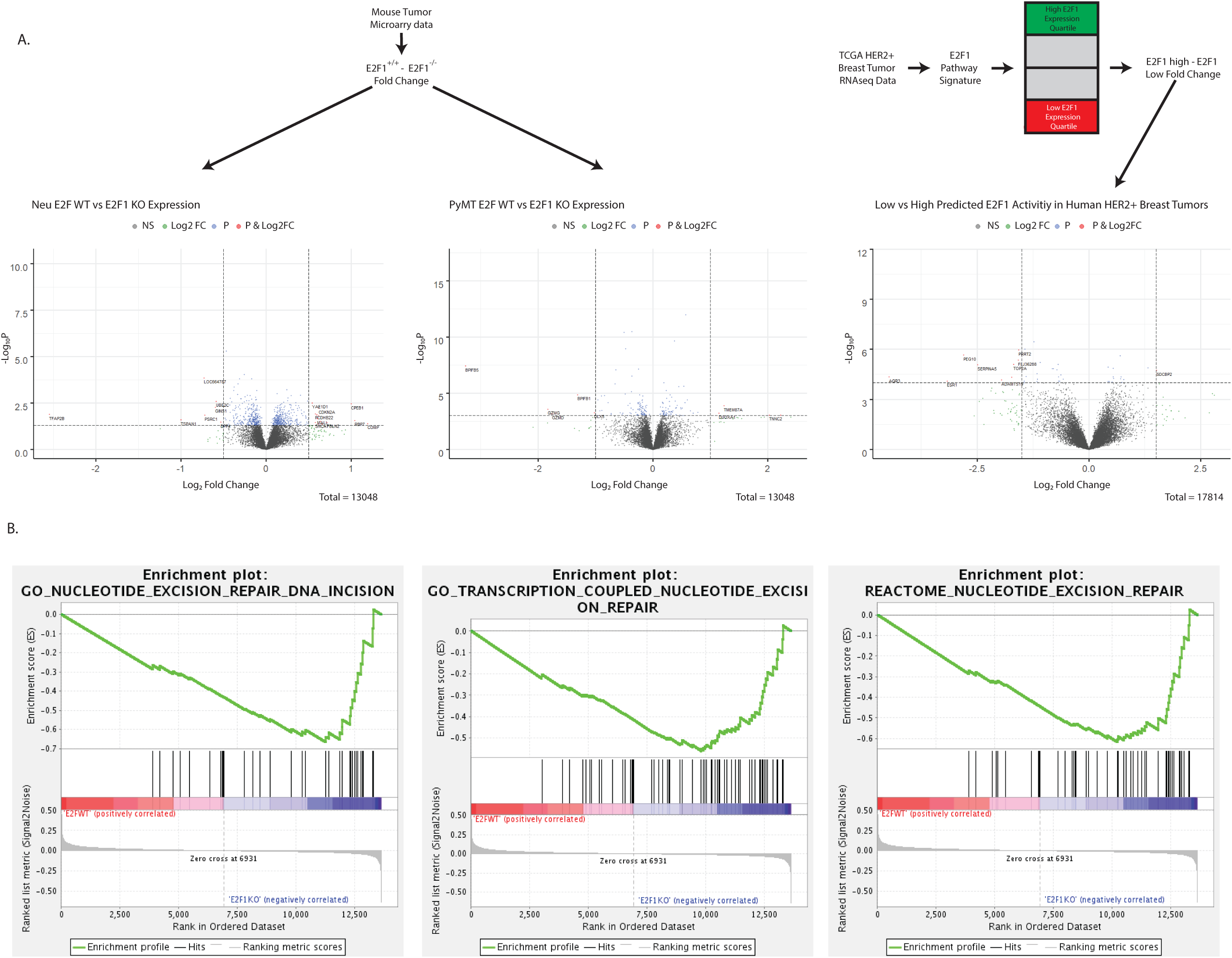
Gene expression changes in E2F1^−/−^ mouse tumors, and E2F1 low human breast cancer. **A)** Two volcano plots show significant fold changes in genes from Neu and PyMT mouse tumors respectively. Fold change was determined by subtracting the E2F1 KO mean from the E2F1 WT mean for each gene. Fold change and p-value cutoff for Neu tumors was .5, and .05 respectively. Fold change and Pvalue for PyMT tumors was 1.0 and .001 respectively. **B)** Diagram represents data processing steps for human TCGA data. A volcano plot shows significant fold change genes in E2F1 high vs. E2F1 low human HER2+ve tumors. Fold change was determined by subtracting samples in the lowest E2F1 quartile mean from the highest E2F1 quartile mean for each gene. Fold change cutoff and p-value for human tumors was 2.0, and 10e^−60^ respectively. C) GSEA plots generated for E2F1 WT vs E2F1 KO tumors (Neu and PyMT combined) show enrichment of Nucleotide excision repair, and WNT signaling pathways in E2F1 KO tumors.

Given that the gene expression analysis did not identify a mechanism altering metastatic potential, we examined genomic events occurring in Neu and PyMT tumors in the E2F1^+/+^ and E2F1^−/−^ backgrounds. Whole genome sequencing was completed and single nucleotide variant profiles were called for each tumor using TCGA best practices. Initial analysis of the single nucleotide variant (SNV) data showed an extremely high proportion of SNVs occurring within chromosome 2 of the E2F1 knockout tumors relative to the E2F1 wild type controls (Figure 3A-B). However, E2F1 is located within the qH1 band of chromosome 2 in mice, and correlates to where the increased proportion of SNVs were observed (Figure 3E). While the E2F1 knockout mice were backcrossed 12 generations to FVB, we hypothesized this abundance of SNVs was called due to residual background strain DNA from the E2F1 knockout stain. Given that E2F1 mice were generated in the SV129 background, and Neu and PyMT mice are on the FVB background, we filtered SNV calls using a list of SNVs that were generated from comparing the SV129 background against the C57/BL6 background, which is the standard mouse reference genome (Figure 3F). As a result, the majority of chromosome 2 SNV calls were filtered out, and the proportion of SNVs was roughly equal across the 19 autosomal mouse chromosomes in our E2F1 WT and E2F1 KO Neu and PyMT tumors (Figure 3G). This serves as an important caution when sequencing mouse models resulting from crosses or not in purebred backgrounds.

**Figure 3:**
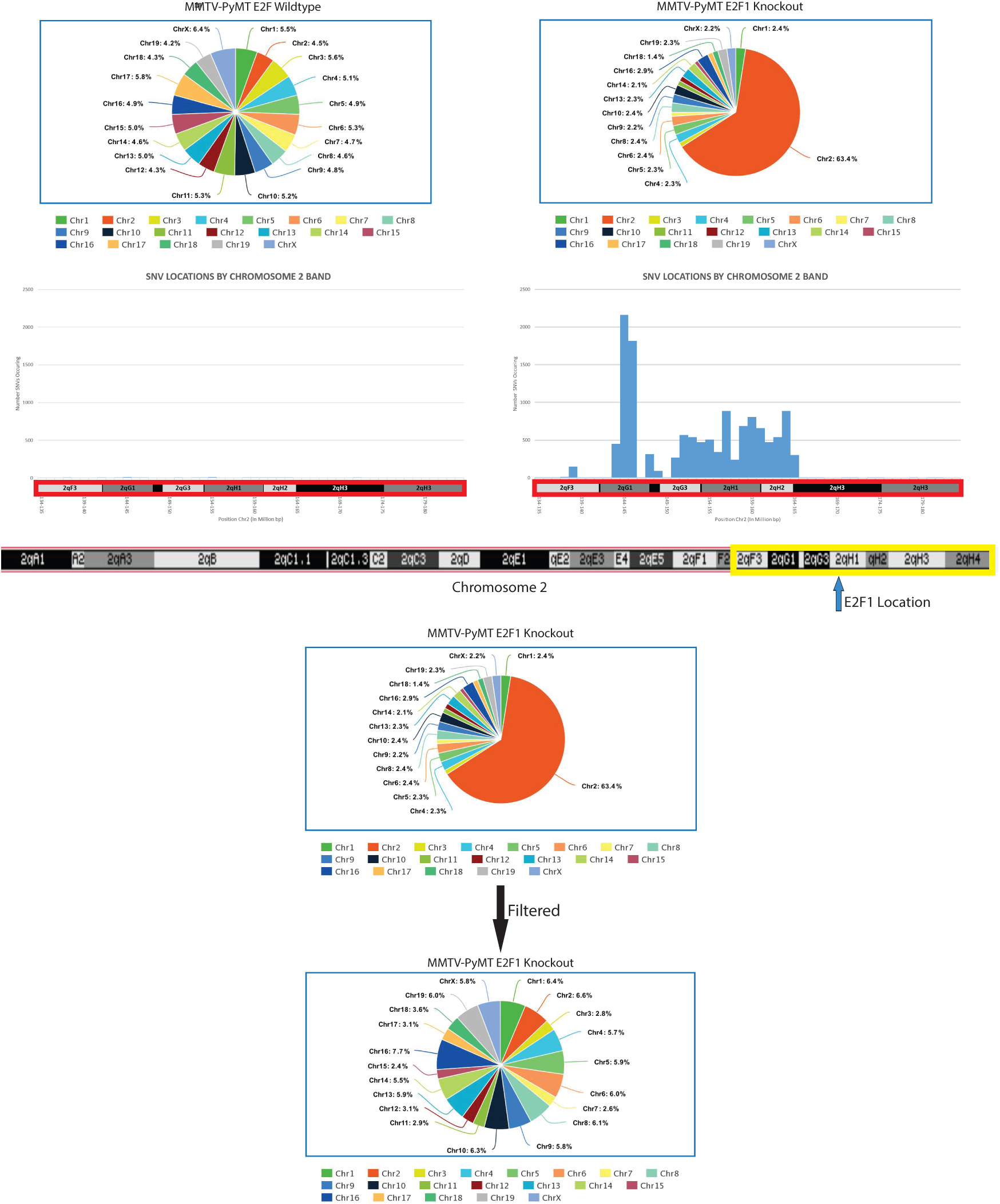
Filtering background strain to remove artifacts that have potential to confound analysis. **A)** Pie chart from an E2F1^+/+^ PyMT tumor represents the normalized (SNVs/Chromosome Size) percentage of SNVs within each chromosome. **B)** Pie chart from an E2F1^−/−^ PyMT tumor represents the normalized percentage of SNVs within each chromosome. An abundance of SNVs within chromosome 2 is observed. **C)** The banding pattern of mouse chromosome 2. The arrow highlights the location of E2F1, and the yellow box represents the bands represented in D and E. **D)** Manhattan plot shows the number of SNVs occurring within the 2qF3-2qH3 bands of chromosome 2, in the E2F1^+/+^ sample from A. **E)** Manhattan plot shows the number of SNVs occurring within the 2qF3-2qH3 bands of chromosome 2, in the E2F1^−/−^ sample from B. **F)** Top pie chart is the same as in B. Bottom pie chart represents the percentage of SNVs across each chromosome of the same sample as above, after filtering on the sv129 background.

Interestingly, the single nucleotide variant mutation burden was higher in PyMT mice as compared to Neu mice (p-value = 0.05), which was surprising due to the brief latency of PyMT tumors (Figure 4A). Except for one PyMT E2F1 knockout tumor, the rate of exonic single nucleotide mutations ranged from .005-.08 mutations per megabase. This mutation rate is similar to previous rates shown for mouse tumors [28], and is lower than the 1 mutation / megabase exonic mutation rate commonly observed in human breast cancer [24].

**Figure 4:**
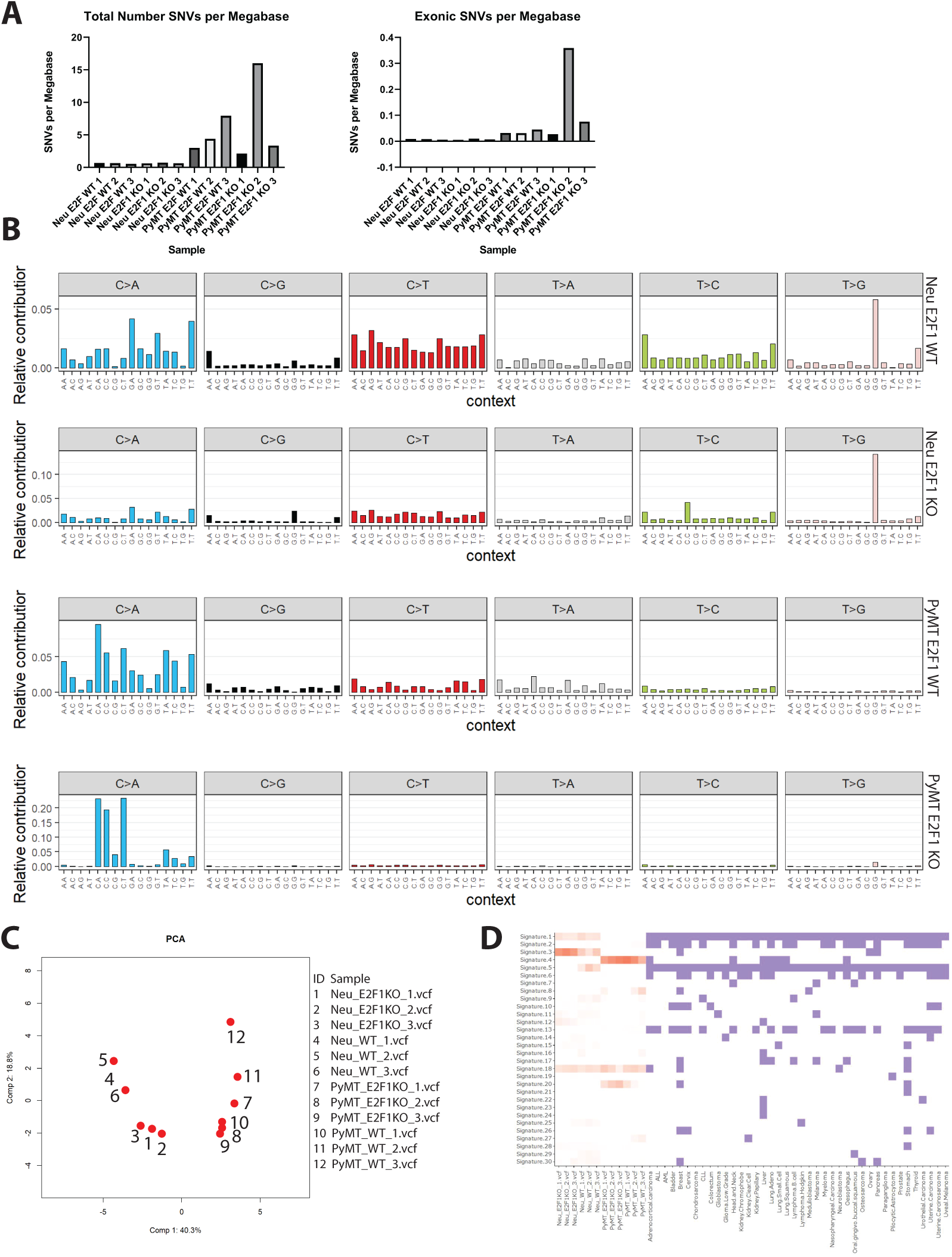
SNV mutation burden in Neu and PyMT tumors. **A)** Bar graphs represent the number of total or exonic mutations per megabase occurring in all 12 sequenced tumors. **B)** Shows representative mutation profiles for each of the four classes of samples sequenced. Mutation profiles are derived from 96 bp trinucleotide signatures originally developed by Alexandrov et. al. Four classes of samples are Neu E2F1^+/+^, Neu E2F1^−/−^, PyMT E2F1^+/+^, PyMT E2F1^−/−^. **C)** PCA plots derived from trinucleotide signatures show clustering of all 12 samples sequenced. **D)** The heatmap of cancer signatures for the 12 sequenced tumors, as well as various cancers is shown.

To analyze distinct types of SNVs occurring within our tumors, and investigate potential mechanisms driving these differences, a mutation signature approach was taken [29]. While trinucleotide signatures showed similarities between Neu and PyMT tumors, there were striking differences, such as T>G mutations occurring almost exclusively in Neu tumors of either E2F1 status (Figure 4B). The signatures for all 12 tumors are shown in Supplemental Figure 1. Principal component analysis (PCA) completed using mutation signatures from all 12 tumors shows distinct clustering between Neu and PyMT tumors (Figure 4C). Furthermore, apart from a single E2F1 KO PyMT tumor, PCA separates E2F1 WT and E2F1 KO tumors into distinct clusters within the Neu and PyMT models.

The contribution of the 30 known COSMIC (catalog of somatic mutations in cancer) signatures to each Neu and PyMT tumor were then determined [29]. While all Neu and PyMT tumors had some contribution from signature 18, there were stark differences in the other COSMIC signatures contributing to Neu and PyMT tumors (Figure 4D). For example, Neu tumors had contributions from signatures 1 and 3, while PyMT tumors were associated with signatures 4 and 20. Furthermore, there were signature differences when comparing E2F1 WT tumors to E2F1 KO tumors within the Neu and PyMT models. For example, Neu E2F1 WT tumors were associated with signatures 5 and 9, while Neu E2F1 KO tumors lacked these associations. When analyzing the proposed etiology for these signatures, Neu tumor signatures are associated with age, while PyMT tumor signatures have no age association, which aligns in the context of Neu and PyMT tumor latency (**supplemental figure 1**). Interestingly, Neu tumors also have an association with inefficient double stranded break repair (DSB), with E2F1 KO tumors being more highly associated than E2F1 WT tumors. PyMT tumor signatures were not associated with DSB, but were highly associated with smoking, and defective DNA mismatch repair (MMR). Together, these data suggest E2F1 loss drives differences in DNA repair and tumor etiology.

Multiple programs were also used to determine copy number variants and translocations occurring within Neu and PyMT tumors (Figure 5A-D). After considering the consensus CNV calls from two programs, over 98% of the copy number events were small in size (under 1 mb), while relatively few larger events (above 1 mb) were observed. When analyzing CNV events from the programs individually, there was a larger proportion of copy number events greater than 1 mb in size. However, these larger events still only accounted for approximately 5% of the copy number events occurring within a tumor. Surprisingly, there was a large amount of copy number gene overlap between the E2F WT and E2F1 KO tumors (Figure 5E). The large number of shared genes involved in copy number events may indicate E2F1 loss is not a primary driver of these events.

**Figure 5:**
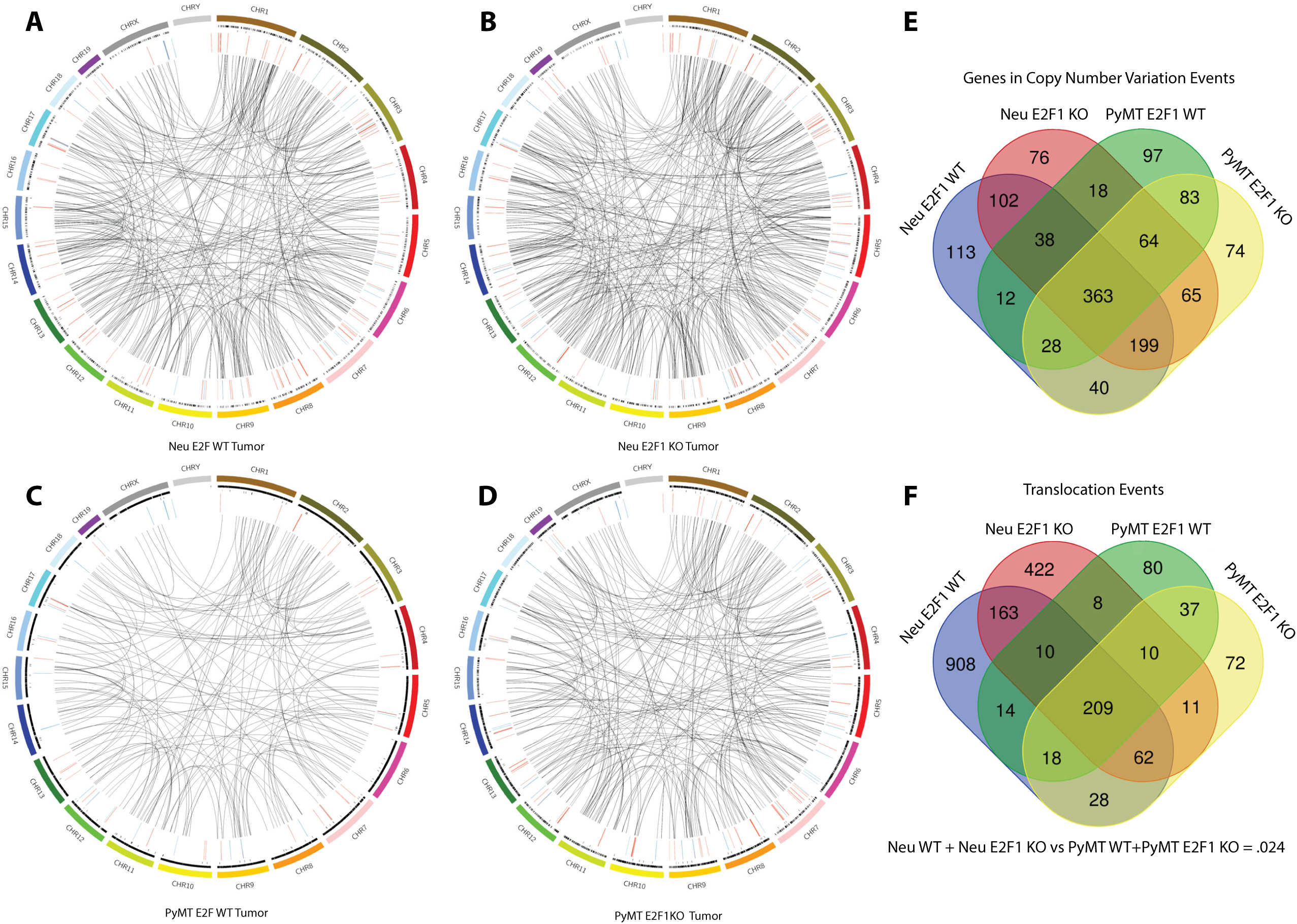
Mutation burden in Neu and PyMT tumors. **A)** Circos plot for a representative Neu E2F1^+/+^ sample. **B)** Circos plot for a representative Neu E2F1^−/−^ sample. **C)** Circos plot for a representative PyMT E2F1^+/+^ sample. **D)** Circos plot for a representative PyMT E2F1^−/−^ sample. For A-D Circos plots, outer most ring represents the mouse chromosomes. Four successive inner rings represent the following mutation types; total SNVs, exonic SNVs, Copy number variation with green being amplification and red being deletion, and translocations. **E)** Venn diagram showing the overlap of genes within copy number events. Consensus copy number events were generated for each of the three samples within the four sample classes. Genes were then extracted and compared across the sample classes. **F)**. Venn diagram showing the overlap of translocations occurring within the four sample classes. Consensus translocations calls from each of the three samples within each class were generated, and the four classes were then compared.

There were a surprisingly large number of translocations occurring within the Neu and PyMT tumors. When comparing the average number of translocations per sample across the genomic models, there were statistically more translocations occurring within Neu tumors than PyMT tumors, regardless of E2F1 status. When comparing across E2F1 status within the each model however, there was no statistically significant difference (Figure 5F). To confirm the translocation calls made by Delly and Lumpy, 20 translocations from each tumor were chosen at random and the read evidence for these translocations was analyzed using Genome Ribbon [43]. Translocation read data for one tumor can be found in Table 1. All tumors had at least 75% of translocations with some read support, with 9 of 12 tumors having at least 85% of translocations with some read support (Supplemental Table 1). Interestingly, all translocation events analyzed had wild type reads present, suggesting a level of heterogeneity occurring within the tumors. To verify one of the translocation events from Table 1, PCR was completed with primers flanking the translocation junction. Both translocated and wild type reads were present at the breakpoint, confirming the existence of the translocation (Figure 6). Based on this evidence, upwards of 80% of the translocations are predicted to be real events.

**Figure 6:**
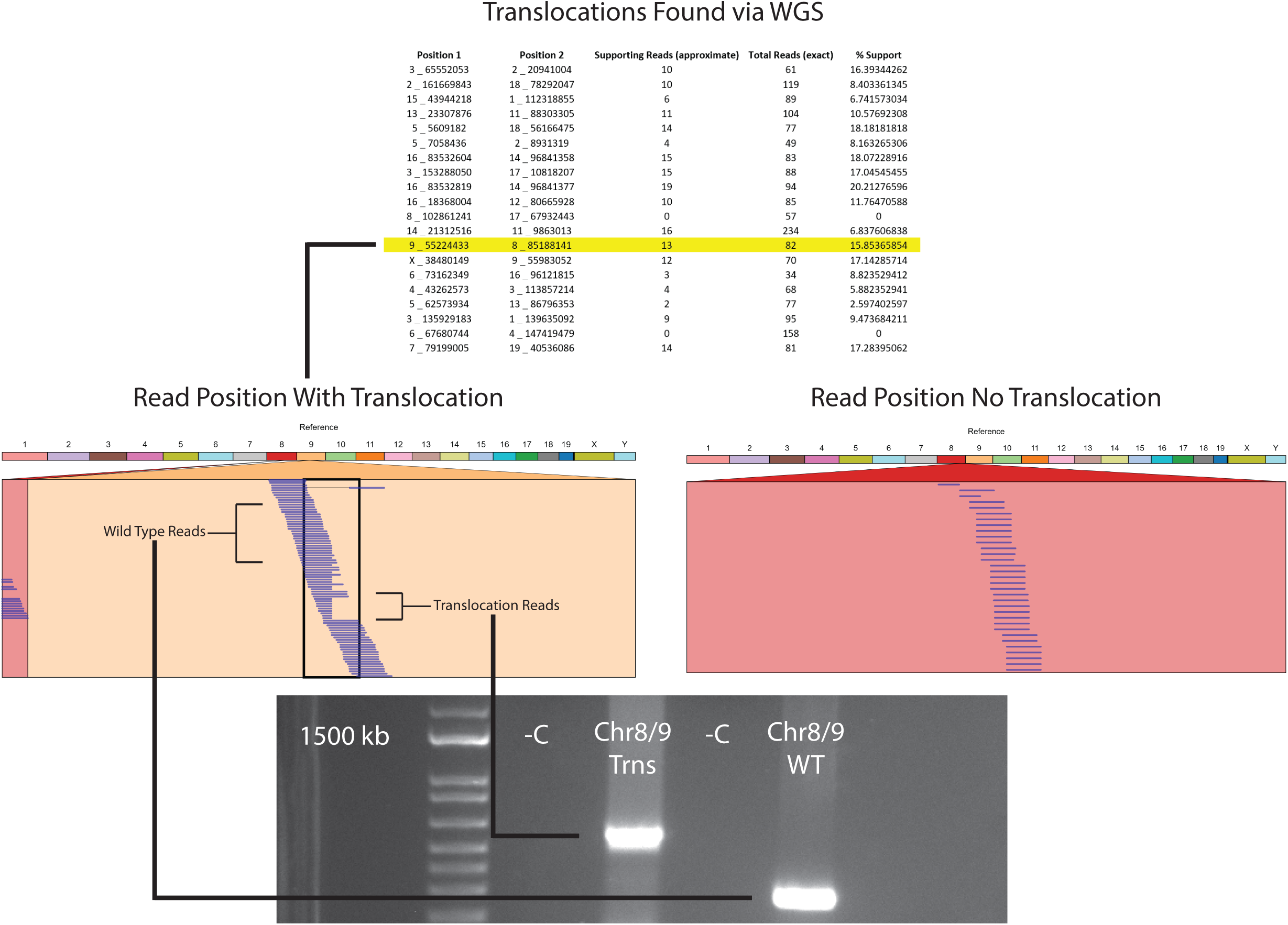
Verification of translocation calls. **A)** Example of a GenomeRibbon plot where no structural variation occurs. The top colored bands represent each chromosome of the mouse, and the red box below represents the location searched within a sample’s bam file. Each line within that box represents a different read. **B)** A GenomeRibbon plot representing translocation number 13 from table 1. Translocated reads are shown between chromosome 9 and chromosome 8. **C)** Gel image of the chromosome 8/9 translocation from the GenomeRibbon plot above. DNA was from a PyMT E2F1^−/−^ tumor. Both translocation and wild type tumor DNA were amplified. Translocated reads were amplified using a primer set flanking the region where the two translocated ends ligate.

To determine whether cancer and metastasis related genes were mutated within E2F1 WT and E2F1 KO tumors, the list of mutations was screened with the list of known cancer genes from the Catalog of Somatic Mutation in Cancer (COSMIC). This analysis found mutations in a number of genes associated with cancer (Supplemental Table 2), but failed to find key metastatic genes consistently mutated within the four sample groups To identify whether an abundance of mutations occurred within particular pathways of E2F1 knockout tumors versus E2F1 wildtype tumors, a database mining approach was taken using Gather [30]. First, genes with potentially impactful mutations were stratified into two gene lists that were distinct in E2F1^−/−^ versus E2F1^+/+^ tumors. Potentially impactful mutations are regarded as SNVs causing stop gain or nonsynonymous mutations, translocations causing truncated or fusion genes, or copy number segments resulting in the amplification or deletion of genes. These two gene lists were then applied to Gather to determine whether Gene Ontology (GO) lists or KEGG pathways were significantly mutated. This analysis determined a number of significant GO lists that were present within the gene list from E2F1^−/−^ tumors, but not E2F1^+/+^ tumors.

One of the most interesting GO lists present within E2F1 mutated tumors, but not E2F1 wildtype tumors, included 48 genes involved in cell adhesion. Many of these genes include various collagens, integrins, and cadherins (Figure 7). Previous research has shown a number of collagens to be important for tumor maintenance, angiogenesis, and metastasis [20]. Collagen IV is the major component of the basement membrane, and is comprised of heterogeneous trimers stemming from six COL4A genes. Three collagen IV genes were found mutated in different PyMT E2F1 KO tumors. Other mutations within PyMT E2F1 KO tumors include COL5A2, with collagen V being a component of the interstitial matrix, COL6A1-3, with collagen VI being abundant in the tumor invasive front [21–23], and various integrin and cadherin genes. Interestingly, when re-analyzing the gene expression data, the integrin pathway was also found dysregulated through Gene Set Enrichment Analysis. There was also an abundance of intronic and synonymous mutations within these genes, suggesting they may be hypermutated due to the disruption of E2F1 within the model. In fact, of the 48 mutated genes from Gather, 24 were predicted to be regulated by E2F1 through TRANSFAC.

**Figure 7:**
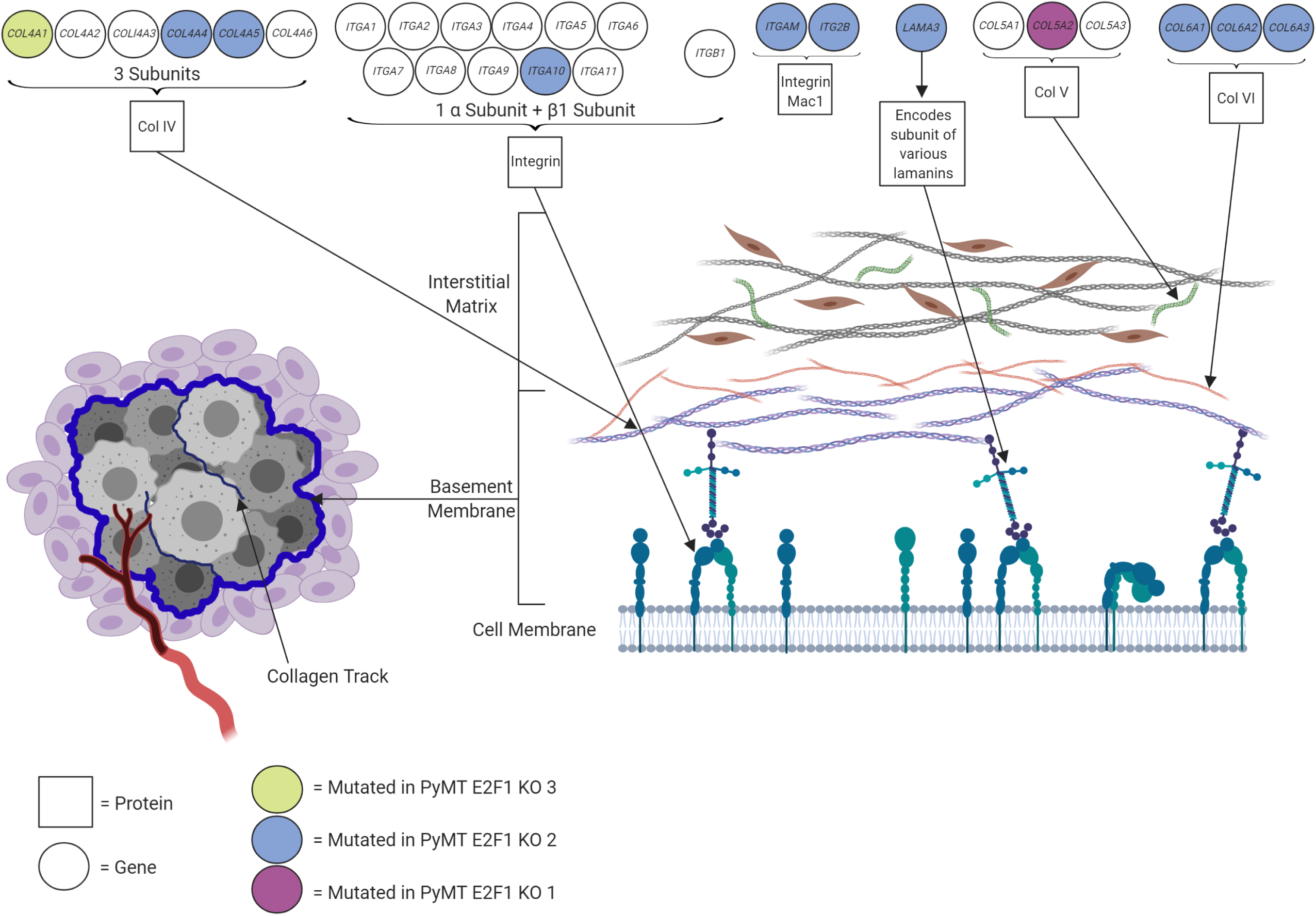
Mutations in basement membrane genes. Diagram shows various mutations occurring in genes that code for proteins making up the basement membrane and interstitial matrix. Circles at top indicate genes with colors representing 1 of 3 sequenced E2F1^−/−^ PyMT tumors that has a mutation in that gene. Image on left represents a breast tumor with surrounding basement membrane. Image on right represents the basement membrane and interstitial matrix on the outer edge of a tumor.

## Discussion

Ablation of E2F1 in PyMT and Neu transgenic mice results in a significant decrease in metastasis to the lung. To determine whether gene expression changes were responsible for the altered phenotypes, microarray data was analyzed but showed no large changes in gene expression between E2F1^+/+^ and E2F1^−/−^ mice. This was recapitulated in human HER2+ breast cancers after separation into E2F1 high/low quartiles. Examination of pathways through GSEA revealed several pathways differentially regulated between E2F1^+/+^ and E2F1^−/−^ tumors, but none of these pathways had obvious implications in regulating metastasis. To test for genomic alterations impacting metastasis, we completed WGS of E2F1^+/+^ and E2F1^−/−^ tumors in both the Neu and PyMT models. Initial analysis showed an abundance of SNV mutations within chromosome 2 of E2F1 knockout mice, but this was corrected by filtering SNVs based on the mouse background. Mutation trinucleotide signatures showed differences between etiology of Neu and PyMT tumors, as well as between the E2F1 knockout and E2F1 WT tumors, with each group clustering separately in a PCA plot. Neu tumors were more closely associated with double stranded break repair, while PyMT tumors were associated with DNA Mismatch Repair. Structural variant analysis showed conservation of copy number genes between the Neu and PyMT models, and a surprisingly high number of translocations within the tumors. Based on the read evidence, we predict greater than 80% of the called translocations are real events. Analyzing the lists of mutated genes for GO and KEGG pathways revealed a significant number of mutated genes involved in cell adhesion. Further analysis of these genes found many of them are involved in the basement membrane and interstitial matrix, which could be a potential route for disruption of the metastatic cascade.

Sequencing data from genetically engineered mouse models is largely lacking, with only a few models having been sequenced [28, 31–33]. SNV mutation rates between previous studies and ours indicate similarities, and the small discrepancies may be explained through differences in data processing methods. Regarding copy number variation, prior research has shown numerous small copy number events, as well as a few larger events [31], although that publication had estimated copy number variation regions from whole exome sequencing data. This was recapitulated in our data, with the exception that large events were not prevalent after taking the consensus of two structural variant callers. We also noted a substantially greater number of translocations within the mouse tumors as compared to a previous study comparing Neu and PyMT E2F1 wildtype tumors, however, the same trend of Neu tumors having more translocations than PyMT tumors holds. This increase in called translocations is likely explained by differences in calling methods, with the previous method being more stringent. Overall, the field may benefit from a large comparison of mouse tumor sequencing data with tumors analyzed under the same parameters.

After analyzing mutated genes using a pathway approach, many genes involved in cell adhesion were found having potentially impactful mutations in E2F1 knockout tumors, but not E2F1 wild type tumors. These included various collagens, integrins, cadherins, and related genes. Of the mutated genes found important to cell adhesion, some, such as *Col4a1*, are important components of the basement membrane and are involved in tumor progression. Disruptions to the basement membrane and other collagen formations has potential to disrupt the metastatic process during pre-intravasation. This theory is supported by previous data we generated, which found a significant decrease in circulating tumor cells [7]. Interestingly, we have also previously noted amplification of *Col1a1* in Neu E2F1 WT tumors, and found this event to be important for the metastatic process [16]. Combined, these data highly suggest various collagens and proteins within the basement membrane are important to the metastatic process in Neu and PyMT tumors.

SNV profiling for human tumors has utility for both discovery and treatment purposes. Sequencing of human breast tumors has revealed larger genomic trends as well as mutation rates for oncogenes and tumor suppressors [34]. Clinically, the benefits of SNV profiling are observed in examples such as EGFR mutant lung cancers being treated with tyrosine kinase inhibitors. The importance of determining SNVs within mouse models is evidenced by previous research from our lab and others [28, 31]. Potential sources of error when determining SNVs can stem from differing genetic background within mice, even after backcrossing, as well as being too loose or too stringent with the filtering process. Interestingly, a prior report identified an important mutation within the *Ptprh* gene [28], but this mutation was not present within this sequence analysis. This discrepancy was due to the different approaches taken for each analysis. While the initial paper stipulated an SNV call must pass 3 of 4 SNV calling programs, the work herein stipulated a call must pass 3 of 3 programs used. When analyzing the SNV data for each program used, a mutation in *Ptprh* was called from SomaticSniper and Varscan, but not called from Mutect2. This suggests the usage of multiple programs to call SNVs is more applicable for discovery purposes, and that less stringent filtering parameters may be beneficial.

When analyzing copy number alterations and translocations within the models, there were a surprising lack of differences across E2F1 status, suggesting E2F1 loss is not a primary driver of these events. The large number of called translocations occurring within both models was surprising, and based on read evidence; we estimate at least 80% of these to be present within the tumors. Furthermore, the varying read support seen for confirmed translocations indicates a high amount of tumor heterogeneity occurring in both models, regardless of E2F1 status. While there were numerous Catalog of Somatic Mutations in Cancer (COSMIC) associated genes mutated within the models, no mutations conserved between E2F1 knockout tumors (within or across models) immediately jumped out as important to the metastatic process.

Analyzing gene expression changes between E2F1 WT and E2F1 KO tumors showed no major changes upon E2F1 loss. This was recapitulated among human HER2+ve breast cancer tumors stratified between low and high E2F1 activity. The lack of major changes on a gene expression level may indicate that compiled small changes are enough to result in phenotypic alterations, or that genomic mutations are leading to altered protein function/localization. Interestingly, the gene encoding Transcription Factor AP-2 Beta was significantly upregulated in Neu E2F1 KO mice. This, combined with the data showing a lack of major gene expression changes between E2F1 WT and E2F1 KO tumors indicates a level of compensation is occurring by other members of the E2F family, as well as other transcription factors [7, 8]. The sequencing data from E2F1^−/−^ Neu and PyMT mice indicate phenotypic changes may be due to an abundance of mutations in particular pathways in addition to minor expression changes observed in microarray data. Taking into consideration that the metastatic process likely originates from a small population of metastatic cells within the primary tumor, the contribution of a few metastatic cells to the bulk tumor gene expression or sequencing data may also cause key events to be lost within the noise of the primary tumor. Future work will address these issues through single cell sequencing and gene expression in matched primary and metastatic tumors.

## Methods

### Gene Expression Analysis

Gene expression data was obtained as described previously [6, 8]. Volcano plots for Neu and PyMT tumors were generated by removing outliers for each sample group using Nowaclean (Holsb, Einar. 2017. “Outlier detection with nowaclean.”), samples that were at least 3.0 standard deviations away when constructing a PCA plot were removed. Data were log2 transformed, and the mean for each gene was calculated within the four sample groups. Fold change was then calculated by subtracting the E2F1 KO mean from the E2F1 WT mean for each gene. P-values were calculated, and data was plotted using EnhancedVolcano (Blighe, Kevin. 2018. “EnhancedVolcano: Publication-ready volcano plots with enhanced colouring and labeling.”) in R. Human RSEM normalized RNAseq breast cancer data from TCGA was downloaded from UCSC Xena, filtered to HER2+ samples, and sorted by E2F1 expression. The lower and upper quartiles were kept, and the data were processed for volcano plots as above. GSEA plots were generated from combining Neu and PyMT gene expression datasets. Datasets were collapsed, and combatted to remove batch effects. GSEA was run using GenePattern [35] from the Broad Institute.

### Whole Genome Sequencing and Processing

Three samples from each group, (for a total of 12), were used to extract DNA from flash frozen tumors according to the manufacture’s protocol of the Qiagen Genomic-tip 20/G kit. Sequencing was completed at a depth of 40x with paired end, 150 base pair reads. DNA was prepped for sequencing using Illumina TruSeq Nano DNA library preparation, and sequenced on an Illumina HiSeq 2500. After sequencing, raw fastq files assessed for quality control using FASTQC (https://www.bioinformatics.babraham.ac.uk/projects/fastqc/), trimmed using Trimmomatic [36], and assessed for quality control for a second time. Files were aligned to the mm10 mouse reference using BWA MEM [37] under standard parameters. After alignment, Picard tools (“Picard Toolkit.” 2019. Broad Institute, GitHub Repository. http://broadinstitute.github.io/picard/; Broad Institute) was used to add read groups, and Samtools [38] was used to sort. PCR duplicates were then removed using Picard tools, and the files were indexed.

### Variant Calling

Somatic SNVs were called using SomaticSniper [39], Mutect2 [40], and VarScan [41]. Consensus calls were merged using R (R Core Team (2018). R: A language and environment for statistical computing. R Foundation for Statistical Computing, Vienna, Austria. Available online at https://www.R-project.org/) base programming, and mutations were only kept if they were called by all three programs. SNV calls were also filtered using base R to account for differences between the FVB strain and mm10 alignment (C57/BL6), as well as differences between the SV129 strain (original E2F1 mouse background) and C57/BL6. SNVs were annotated using Annovar [42]. CNVs were determined by keeping the consensus of Lumpy [43] and Delly [44]. Consensus was determined using Intansv (Yao W (2019). intansv: Integrative analysis of structural variations. R package version 1.24.0) at a threshold of .2, and events smaller than 10,000 bp were filtered out. Intansv was also used to annotate CNV events. Translocations were called using Lumpy and Delly, and filtered based on read evidence. Lumpy calls were kept if they has at least 20 supporting split end and paired end reads, Delly calls were kept if there was split end and paired end read evidence for the call. Where needed, WT FVB mouse sequence was used as a normal control.

### Mutation Signatures

Trinucleotide mutation signatures were completed for each sample using the Musica [45] shiny app in R. Musica code was altered to allow for the use of the mouse mm10 reference genome.

### Circos plots

Circos plots were generated for each sample using CIRCOS version .69 [46]. Genetic variants were plotted according to the mm10 reference genome.

### Translocation Verification

Read evidence for 20 randomly selected translocations from all 12 sequenced samples was examined using GenomeRibbon [47]. For PCR verification, primers were designed with at least 400 bp flanking the predicted breakpoint.

## Acknowledgements

This work was supported with NIH R01CA160514 and Worldwide Cancer Research WCR - 14-1153.

## References

1. Guy CT, Webster MA, Schaller M, Parsons TJ, Cardiff RD, Muller WJ (1992) Expression of the neu protooncogene in the mammary epithelium of transgenic mice induces metastatic disease. Proc Natl Acad Sci 89:10578–10582

2. Guy CT, Cardiff RD, Muller WJ (1992) Induction of mammary tumors by expression of polyomavirus middle T oncogene: a transgenic mouse model for metastatic disease. Mol Cell Biol 12:954–961

3. Cardiff RD, Anver MR, Gusterson B a, et al (2000) The mammary pathology of genetically engineered mice: the consensus report and recommendations from the Annapolis meeting. Oncogene 19:968–988

4. Herschkowitz JI, Simin K, Weigman VJ, et al (2007) Identification of conserved gene expression features between murine mammary carcinoma models and human breast tumors. Genome Biol 8:R76

5. Hollern DP, Andrechek ER (2014) A genomic analysis of mouse models of breast cancer reveals molecular features of mouse models and relationships to human breast cancer. Breast Cancer Res. doi: 10.1186/bcr3672

6. Hollern DP, Swiatnicki MR, Andrechek ER (2018) Histological subtypes of mouse mammary tumors reveal conserved relationships to human cancers. PLoS Genet. doi: 10.1371/journal.pgen.1007135

7. Hollern DP, Honeysett J, Cardiff RD, Andrechek ER (2014) The E2F Transcription Factors Regulate Tumor Development and Metastasis in a Mouse Model of Metastatic Breast Cancer. Mol Cell Biol 34:3229–3243

8. Andrechek ER (2015) HER2/Neu tumorigenesis and metastasis is regulated by E2F activator transcription factors. Oncogene. doi: 10.1038/onc.2013.540

9. Nevins JR (1992) E2F: A link between the Rb tumor suppressor protein and viral oncoproteins. Science (80-) 258:424–429

10. Gorgoulis VG, Zacharatos P, Mariatos G, Kotsinas A, Bouda M, Kletsas D, Asimacopoulos PJ, Agnantis N, Kittas C, Papavassiliou AG (2002) Transcription factor E2F-1 acts as a growth-promoting factor and is associated with adverse prognosis in nonsmall cell lung carcinomas. J Pathol 198:142–156

11. Qin G, Kishore R, Dolan CM, et al (2006) Cell cycle regulator E2F1 modulates angiogenesis via p53-dependent transcriptional control of VEGF.

12. Rouaud F, Hamouda-Tekaya N, Cerezo M, et al (2018) E2F1 inhibition mediates cell death of metastatic melanoma. Cell Death Dis. doi: 10.1038/s41419-018-0566-1

13. Field SJ, Tsai FY, Kuo F, Zubiaga AM, Kaelin WG, Livingston DM, Orkin SH, Greenberg ME (1996) E2F-1 Functions in mice to promote apoptosis and suppress proliferation. Cell 85:549–561

14. Cancer Genome Atlas Research Network JN, Weinstein JN, Collisson EA, Mills GB, Shaw KRM, Ozenberger BA, Ellrott K, Shmulevich I, Sander C, Stuart JM (2013) The Cancer Genome Atlas Pan-Cancer analysis project. Nat Genet 45:1113–20

15. Chaffer CL, Weinberg RA (2011) A perspective on cancer cell metastasis. Science (80-) 331:1559–1564

16. Fidler IJ (2003) The pathogenesis of cancer metastasis: The “seed and soil” hypothesis revisited. Nat Rev Cancer 3:453–458

17. Welch DR, Hurst DR (2019) Defining the Hallmarks of Metastasis. Cancer Res 79:3011–3027

18. Burnier J V., Wang N, Michel RP, et al (2011) Type IV collagen-initiated signals provide survival and growth cues required for liver metastasis. Oncogene 30:3766–3783

19. Chang TT, Thakar D, Weaver VM (2017) Force-dependent breaching of the basement membrane. Matrix Biol 57–58:178–189

20. Walker C, Mojares E, del Río Hernández A (2018) Role of Extracellular Matrix in Development and Cancer Progression. Int J Mol Sci 19:3028

21. Cox TR, Rumney RMH, Schoof EM, et al (2015) The hypoxic cancer secretome induces pre-metastatic bone lesions through lysyl oxidase. Nature 522:106–110

22. Daves MH, Hilsenbeck SG, Lau CC, Man TK (2011) Meta-analysis of multiple microarray datasets reveals a common gene signature of metastasis in solid tumors. BMC Med Genomics. doi: 10.1186/1755-8794-4-56

23. Cosphiadi I, Atmakusumah TD, Siregar NC, Muthalib A, Harahap A, Mansyur M (2018) Bone Metastasis in Advanced Breast Cancer: Analysis of Gene Expression Microarray. Clin Breast Cancer 18:e1117–e1122

24. Lawrence MS, Stojanov P, Polak P, et al (2013) Mutational heterogeneity in cancer and the search for new cancer-associated genes. Nature 499:214–218

25. Ding L, Ellis MJ, Li S, et al (2010) Genome remodelling in a basal-like breast cancer metastasis and xenograft. Nature 464:999–1005

26. Yachida S, Jones S, Bozic I, et al (2010) Distant metastasis occurs late during the genetic evolution of pancreatic cancer. Nature 467:1114–1117

27. Navin N, Kendall J, Troge J, et al (2011) Tumor evolution inferred by single-cell sequencing. Nature 472:90–95

28. Rennhack JP, To B, Swiatnicki M, et al (2019) Integrated analyses of murine breast cancer models reveal critical parallels with human disease. Nat Commun 10:3261

29. Alexandrov LB, Nik-Zainal S, Wedge DC, et al (2013) Signatures of mutational processes in human cancer. Nature 500:415–421

30. Chang JT, Nevins JR (2006) GATHER: a systems approach to interpreting genomic signatures. Bioinformatics 22:2926–2933

31. McFadden DG, Politi K, Bhutkar A, et al (2016) Mutational landscape of EGFR-, MYC-, and Kras-driven genetically engineered mouse models of lung adenocarcinoma. Proc Natl Acad Sci 113:E6409–E6417

32. Francis JC, Melchor L, Campbell J, et al (2015) Whole-exome DNA sequence analysis of Brca2 – And Trp53-deficient mouse mammary gland tumours. J Pathol 236:186–200

33. Campbell KM, O’Leary KA, Rugowski DE, Mulligan WA, Barnell EK, Skidmore ZL, Krysiak K, Griffith M, Schuler LA, Griffith OL (2019) A Spontaneous Aggressive ERα+ Mammary Tumor Model Is Driven by Kras Activation. Cell Rep 28:1526–1537.e4

34. Shah SP, Roth A, Goya R, et al (2012) The clonal and mutational evolution spectrum of primary triple-negative breast cancers. Nature 486:395–399

35. Reich M, Liefeld T, Gould J, Lerner J, Tamayo P, Mesirov JP (2006) GenePattern 2.0. Nat Genet 38:500–501

36. Bolger AM, Lohse M, Usadel B (2014) Trimmomatic: a flexible trimmer for Illumina sequence data. Bioinformatics 30:2114–2120

37. Li H (2013) Aligning sequence reads, clone sequences and assembly contigs with BWA-MEM.

38. Li H, Handsaker B, Wysoker A, Fennell T, Ruan J, Homer N, Marth G, Abecasis G, Durbin R (2009) The Sequence Alignment/Map format and SAMtools. Bioinformatics 25:2078–2079

39. Larson DE, Harris CC, Chen K, Koboldt DC, Abbott TE, Dooling DJ, Ley TJ, Mardis ER, Wilson RK, Ding L (2012) Somaticsniper: Identification of somatic point mutations in whole genome sequencing data. Bioinformatics 28:311–317

40. Cibulskis K, Lawrence MS, Carter SL, Sivachenko A, Jaffe D, Sougnez C, Gabriel S, Meyerson M, Lander ES, Getz G (2013) Sensitive detection of somatic point mutations in impure and heterogeneous cancer samples. Nat Biotechnol. doi: 10.1038/nbt.2514

41. Koboldt DC, Zhang Q, Larson DE, Shen D, McLellan MD, Lin L, Miller CA, Mardis ER, Ding L, Wilson RK (2012) VarScan 2: Somatic mutation and copy number alteration discovery in cancer by exome sequencing. Genome Res 22:568–576

42. Wang K, Li M, Hakonarson H (2010) ANNOVAR: Functional annotation of genetic variants from high-throughput sequencing data. Nucleic Acids Res. doi: 10.1093/nar/gkq603

43. Layer RM, Chiang C, Quinlan AR, Hall IM (2014) LUMPY: A probabilistic framework for structural variant discovery. Genome Biol 15:R84

44. Rausch T, Zichner T, Schlattl A, Stütz AM, Benes V, Korbel JO (2012) DELLY: Structural variant discovery by integrated paired-end and split-read analysis. Bioinformatics 28:333–339

45. Díaz-Gay M, Vila-Casadesús M, Franch-Expósito S, Hernández-Illán E, Lozano JJ, Castellví-Bel S (2018) Mutational Signatures in Cancer (MuSiCa): A web application to implement mutational signatures analysis in cancer samples. BMC Bioinformatics 19:224

46. Krzywinski M, Schein J, Birol I, Connors J, Gascoyne R, Horsman D, Jones SJ, Marra MA (2009) Circos: An information aesthetic for comparative genomics. Genome Res 19:1639–1645

47. Nattestad M, Chin C-S, Schatz MC (2016) Ribbon: Visualizing complex genome alignments and structural variation. bioRxiv 0344:82123

